# Accelerated dystrophy and decay of oligodendrocyte precursor cells in the APP/PS1 model of Alzheimer’s-like pathology

**DOI:** 10.1101/2020.09.23.309666

**Authors:** Irene Chacon-De-La-Rocha, Gemma Fryatt, Andrea Rivera, Alex Verkhratsky, Olivier Raineteau, Diego Gomez-Nicola, Arthur M. Butt

## Abstract

Myelin disruption is a feature of natural aging and of Alzheimer’s disease (AD). In the CNS, myelin is produced by oligodendrocytes, which are generated throughout life by oligodendrocyte progenitor cells (OPCs). Here, we examined age-related changes in OPCs in APP/PS1 mice, a model for AD-like pathology, compared with non-transgenic (Tg) age-matched controls. Analysis was performed in the CA1 area of the hippocampus following immunolabelling for NG2 with the nuclear dye Hoescht, to identify OPC and OPC sister cells, a measure of OPC replication, together with Gpr17 and Olig2 for oligodendrocytes and myelin basic protein (MBP) immunostaining as a measure of myelination. The results indicate a decrease in the number of OPCs between 9 and 14 months in natural ageing and this occurred earlier at 9 months in APP/PS1 mice, without further decline at 14 months. The number of OPC sister cells was unaltered in natural aging, but declined significantly at 14-months in APP/PS1 mice. The number of GPR17+ and Olig2+ oligodendrocytes was not altered in APP/PS1, whereas MBP immunostaining increased between 9 and 14 months in natural ageing, but not in APP/PS1 mice. Notably, OPCs displayed marked morphological changes at 14 months in APP/PS1 mice, characterized by an overall shrinkage of OPC process domains and increased process branching, characteristic of reactive pathological changes. The results indicate that OPC and myelin disruption are pathological signs in the APP/PS1 mouse model of AD.

## Introduction

Alzheimer’s disease (AD) is the most common type of dementia and it is characterized by the formation of intracellular neurofibrillary tangles (NFTs) and extracellular amyloid-β (Aβ) plaques (Braak and Braak, 1991). White matter disruption is present at an early stage of AD pathology (Bartzokis, 2011, Ihara et al., 2010), and post-mortem analyses indicate that a loss of oligodendrocytes in AD could serve as a diagnostic tool for differentiating white matter pathologies in dementia (Sjöbeck and Englund, 2003, Brickman et al., 2015). Studies in human AD and mouse models indicate loss of oligodendrocytes and demyelination is most pronounced at the core of Aβ plaques (Mitew et al., 2010). Hence, myelin loss is a feature of human AD and mouse models (Desai et al., 2009), but the underlying causes are unresolved.

In the adult brain, oligodendrocyte progenitor cells (OPCs) are responsible for the life-long generation of oligodendrocytes required to myelinate new connections, formed in response to new life experiences, and to replace myelin lost in pathology (Xiao et al., 2016, McKenzie et al., 2014, Young et al., 2013, Hughes et al., 2018). OPCs are identified by their expression of the NG2 proteoglycan and are sometimes known as NG2-cells or NG2-glia (Butt et al., 2002). Prior to differentiating into mature myelinating oligodendrocytes, OPCs transition through an intermediate phase identified by expression of the G-protein coupled receptor GPR17 (Viganò et al., 2016). Notably, early changes in OPCs may be a pathological sign and underlie myelin loss in mouse models of AD-like pathology (Rivera et al., 2016, Vanzulli et al., 2020). This possibility is supported by immunostaining of post-mortem AD brain showing reduced NG2 immunoreactivity in individuals with high Aβ plaque load (Nielsen et al., 2013b).

The APP/PS1 transgenic mouse expresses familial AD-causing mutated forms of human APP (APPswe, Swedish familial AD-causing mutation) and presenilin1 (PS1dE9) and is used extensively as a model for AD-like pathology (Borchelt et al., 1997). The APP/PS1 mouse presents early Aβ plaque deposition in the hippocampus at 4-5 months of age and extensively throughout the forebrain by 8 months (Borchelt et al., 1997). Histological, gene expression and imaging analyses indicate white matter and myelin disruption in APP/PS1 mice (Dong et al., 2018, Wu et al., 2017, Shu et al., 2013). However, the age-related changes in OPCs in APP/PS1 are unclear, with one study indicating an early decrease in NG2 cells in white matter (Dong et al., 2018), and another indicating increased NG2 cells clustering around Aβ plaques (Kulijewicz-Nawrot et al., 2013). Here, we examined age-related changes in OPCs and oligodendrocytes in APP/PS1 mice compared to age-matched non-transgenic controls and focused on the AD-relevant CA1 area of the hippocampus, where disruption of myelination is a prominent feature of AD in humans and the APP/PS1 model (Ota et al., 2019, Chao et al., 2018). Our results indicate an age-related decline in OPC density was accelerated in APP/PS1 and occurred at 9 months compared to 14 months in natural ageing. Notably, OPCs displayed cellular shrinkage and increased process branching, characteristic of reactive changes in response to pathology (Ong and Levine, 1999). These changes in OPCs at 14 months corresponded to a decline in OPC sister cells, a measure of OPC self-renewal, and decreased MBP immunostaining, although overall numbers of GPR17+ and Olig2+ oligodendrocytes were not significantly changed. This study identifies pathological changes in OPCs in the APP/PS1 mouse model of AD and allow a more refined understanding of the progressive changes associated to myelin loss.

## MATERIAL AND METHODS

### Animals and tissue

All procedures were carried out in accordance with the Animals (Scientific Procedures) Act 1986 of the UK. Transgenic APP/PS1 mice were used contain human transgenes for both APP (KM670/671NL, Swedish) and PSEN1 (L166P) (Radde et al., 2006, Maia et al., 2013). APP/PS1 mice and age matched non-transgenic controls aged 9 and 14 months perfusion fixed intracardially under terminal anaesthesia with 4% paraformaldehyde (PFA), then post-fixed for 2 hours with 4% PFA. Sections were cut on a vibratome (Leica) at a thickness of 35 μm then stored in cryoprotectant at −70°C until use.

### Immunohistochemistry

Sections were treated for a blocking stage of either 10-20% normal goat serum (NGS) or normal donkey serum (NDS) or 0.5% bovine serum albumin (BSA) for 1-2 h, depending on the primary antibodies to be used. Sections were washed 3 times in PBS, and incubated overnight in primary antibody diluted in blocking solution containing 0.25% Triton-X: rabbit anti-NG2, 1:500 (Millipore); rabbit anti-Olig2, 1:500 (Millipore); rabbit anti-GPR17, 1:100 (Cayman Labs); rat anti-MBP, 1:300 (Millipore). Sections were washed 3 times in PBS, and incubated overnight in primary antibody diluted in blocking solution containing 0.25% Triton-X: rabbit anti-NG2, 1:500 (Millipore); rabbit anti-Olig2, 1:500 (Millipore); rabbit anti-GPR17, 1:100 (Cayman Labs); rat anti-MBP, 1:300 (Millipore). Tissues were then washed 3 times in PBS and incubated with appropriate fluorochrome secondary antibody (AlexaFluor^®^ 488, AlexaFluor^®^ 568, 1:400, Life Technologies), or biotinylated secondary antibody (Vector Labs) diluted in blocking solution for 1-2h. Finally, sections were washed 3 times with PBS before being mounted on glass slides and covered with mounting medium and glass coverslips ready for imaging.

### Imaging and Analysis

Immunofluoresecence images were captured using a Zeiss Axiovert LSM 710 VIS40S confocal microscope and maintaining the acquisition parameters constant to allow comparison between samples within the same experiment. Acquisition of images for cell counts was done with x20 objective. Images for OPC reconstruction were taken using x100 objective and capturing z-stacks formed by 80-100 single plains with an interval of 0.3 μm. Cell counts were performed in a constant field of view (FOV) of 100 μm x100 μm or 200 μm x 200 μm, depending on the area analysed, in projected flattened images from z-stacks formed by 10 or 15 *z*-single plain images with 1μm interval between them. The relative density of MBP immunolabelling was measured within a constant FOV using ImageJ. For DAB immunostaining of Olig2+ oligodendrocytes, sections were examined on an Olympus dotSlide digital slide scanning system based on a BX51 microscope stand with integrated scanning stage and Olympus CC12 colour camera. The cell coverage of OPCs was measured using ImageJ by drawing a line around the cell processes and measuring the area enclosed within the line. For morphological analysis, single OPCs were drawn in detail using Neurolucida 360 and their morphology was analysed using Neurolucida 360 explorer. For Sholl analysis, the interval between Sholl shells was 5μm. All data were expressed as Mean±SEM and tested for significance by unpaired t-tests or Mann-Whitney tests as appropriate using GraphPad Prism 6.0.

## RESULTS

### Accelerated decline of OPCs in the hippocampus of APP/PS1 mice

The hippocampus displays a high degree of adult oligodendrogenesis, which is important for learning and plasticity (Steadman et al., 2020). Here, we used NG2 immunolabelling to identify adult OPCs (Nishiyama et al., 2016) in the CA1 area of the hippocampus. The results demonstrate that OPCs are uniformly distributed throughout the hippocampus at both 9 and 14 months, in APP/PS1 mice and age-matched controls (Fig. 1A, B). NG2+ OPCs are often observed as duplets or triplets of recently divided sister cells (some indicated by arrows in Fig. 1A, B), confirming previous studies that adult OPCs are proliferating cells (Young et al., 2013, Psachoulia et al., 2009). Quantification confirmed a 30% decrease in the density of NG2+ OPCs between 9- and 14-months in non-transgenic controls (Fig. 1C; *p*<0.05, unpaired t-test, *n*=3 for each group). Notably, at 9-months in APP/PS1 mice there was a significant 50% decrease in NG2+ OPCs compared to age-matched controls (Fig. 1C; *p*<0.05, unpaired t-test, *n*=3), reaching an OPC density equivalent to that observed at 14 months in natural ageing (Fig. 1C); there was no further decline in OPC numbers between 9 and 14 months APP/PS1 mice, which were the same as age-matched controls (Fig. 1C). In addition, there was a decrease in the number of OPC sister cells in APP/PS1 mice, although this only reached statistical significance at 14 months (Fig. 1D; *p*<0.05, unpaired t-test, *n*=3). The results indicate an age-related decline in OPC numbers is accelerated in APP/PS1 mice, which is not offset by OPC self-renewal.

**Figure 1.**
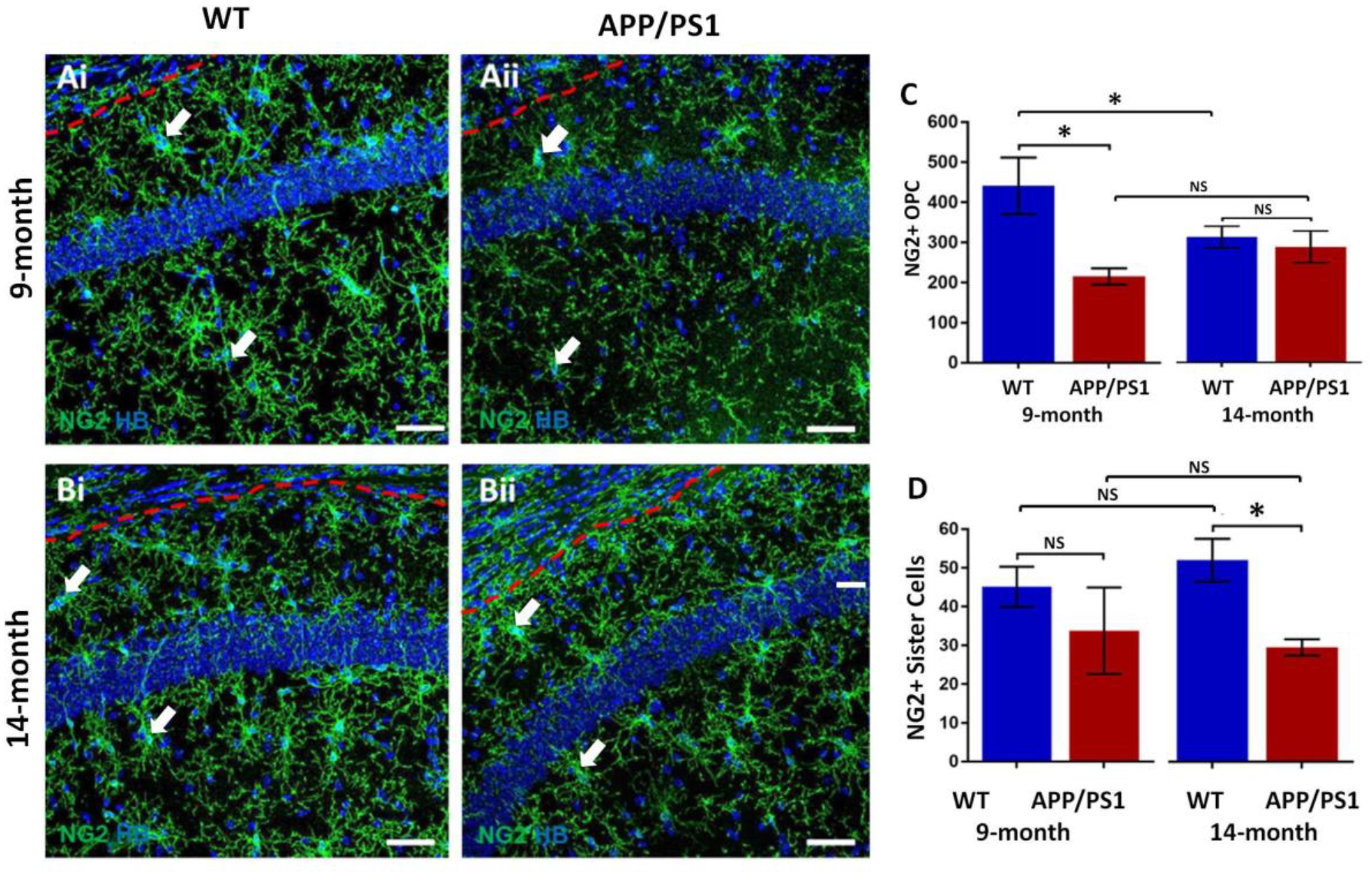
Changes in OPCs in the CA1 area of the hippocampus of APP/PS1 mice. Hippocampi of 9months old and 14-months old APP/PS1 mice were compared to age-matched controls. (**A, B**) Representative confocal images of immunofluorescence labelling for NG2 (green) to identify OPCs and counterstaining with Hoechst (blue) for nuclei, to identify OPC sister cells (some indicated by arrows), from non-transgenic controls (Ai, Bi) and APP/PS1 mice (Aii, Bii), aged 9 months (Ai, Aii) and 14 months (Bi, Bii); scale bars = 50μm. (**C, D**) Bar graphs of total numbers of NG2^+^ OPCs (C) and OPC sister cells (D). Data are expressed as Mean ± SEM; NS= no significant, *p<0,05; Unpaired t-tests, *n*= 3 animals for each group.

### Decline in myelination in the hippocampus of APP/PS1 mice

The hippocampus displays a high degree of myelination, which is essential for cognitive function (Abraham et al., 2010), and myelination has been shown to be disrupted in this area in APP/PS1 mice and it is relevant to AD pathology (Ota et al., 2019, Chao et al., 2018, Dong et al., 2018). We used immunolabelling for MBP to examine myelination (Chao et al., 2018), GPR17 to identify immature oligodendrocytes (Viganò et al., 2016), and Olig2 for oligodendrocytes (Rivers et al., 2008). MBP immunostaining is prominent in the CA1 area at both 9- and 14-months in controls and in APP/PS1 (Fig. 2A, B), as are GPR17+ immature oligodendrocytes, with their characteristic process-bearing morphology (upper insets, Fig. 2A, B), and Olig2 nuclear stained oligodendrocytes (lower insets, Fig. 2A, B). No significant age-related changes in controls or APP/PS1 were observed in the density of GPR17+ and Olig2+ oligodendrocytes (Fig. 2C, D). Measurements of MBP immunostaining density indicated an increase in the CA1 region between 9 and 14 months in non-transgenic controls, consistent with continued myelination up to 15 months of age (Hill et al., 2018), and this was not observed in APP/PS1 mice (Fig. 2E; *p*<0.05, unpaired t-tests, n=3 for both ages). Overall, the results indicate MBP immunostaining is retarded at later stages of pathology in APP/PS1 mice, but the overall numbers of oligodendrocytes are not significantly altered.

**Figure 2.**
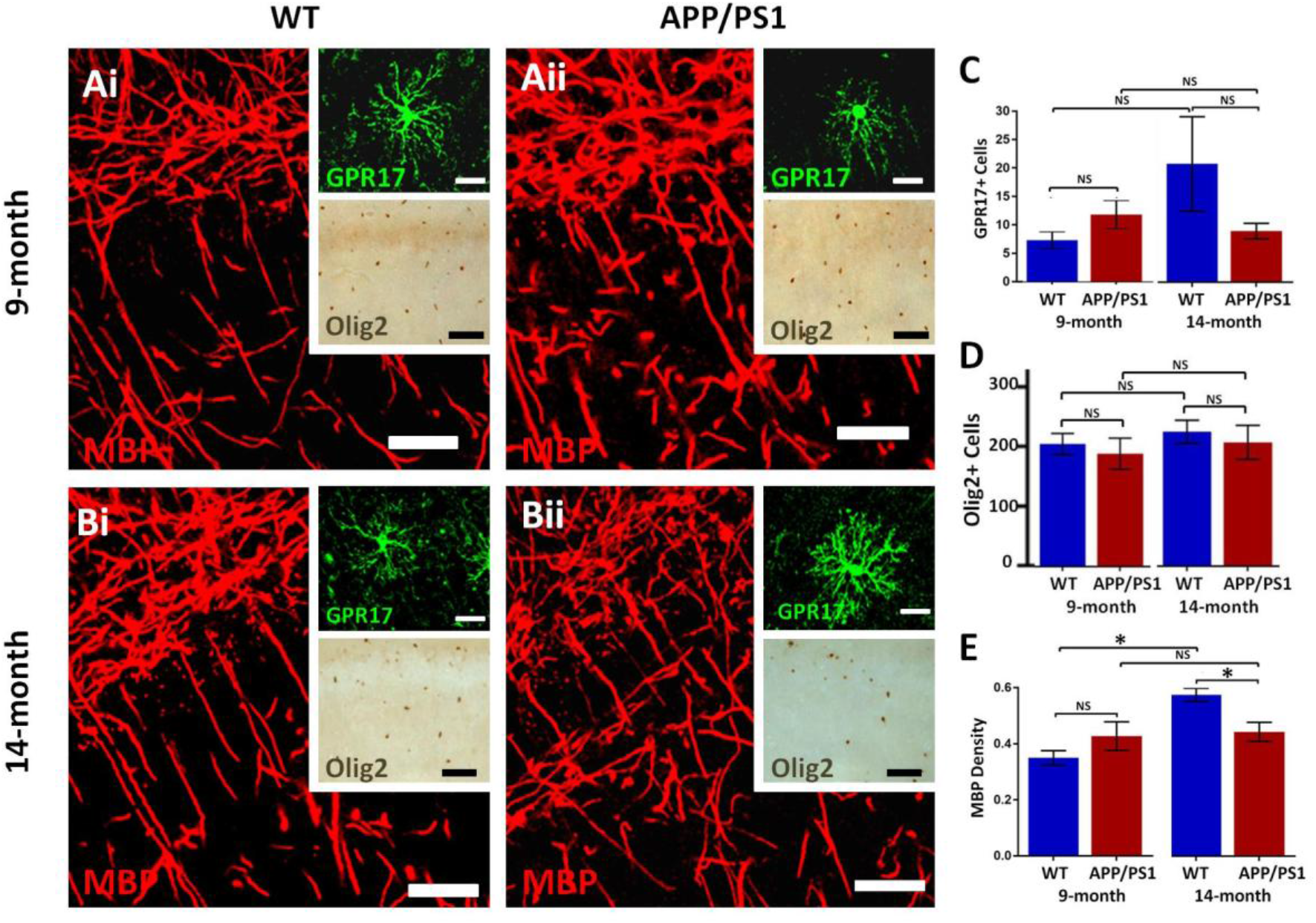
Changes in oligodendrocytes and myelin in the CA1 area of the hippocampus of APP/PS1 mice. Hippocampi of 9months old and 14months old APP/PS1 mice were compared to age-matched controls. (**A, B**) Representative photomicrographs of immunolabelling for MBP (red) to identify the extent of myelination, together with GPR17 for immature oligodendrocytes (upper insets, green) and Olig2 for total number of oligodenrocyte lineage cells (lower panels, brown); scale bars = 50μm, except upper insets = 20 mm. (**C-E**) Bar graphs of numbers of GPR17+ cells (C) and Olig2+ cells (D), together with MBP immunofluorescence density (E); data are expressed as Mean ± SEM, NS= no significant, *p<0,05; Unpaired t-tests, *n*= 3 animals for each group.

### OPC exhibit cellular shrinkage in APP/PS1 mice

The results above indicate OPC are disrupted in APP/PS1 mice, which is often associated with changes in OPC morphology in AD and other pathologies (Butt et al., 2019a, Butt et al., 2019b). We therefore examined OPC morphology in depth, using high magnification confocal images and measuring the process domains of individual cells and the total coverage of NG2 cells within the CA1 (Fig. 3). No differences were observed in OPCs at 9-months in APP/PS1 compared to controls (Fig. 3Ai-Aiv; unpaired t-test, *n*=12 cells per group), whereas at 14-months there was a significant shrinkage of process domains in individual OPCs in APP/PS1 compared to controls (Fig. 3Bi-Biii; *p* < 0.001, unpaired t-test, *n*=12 cells per group). Similarly, overall OPC process coverage in the CA1, measured as NG2 immunostaining within a constant FOV, was significantly decreased at 14-months in APP/PS1 compared to controls (Fig. 3Aiv, Biv; *p*<0.05, unpaired t-test, *n*=3 animals per group). The results indicate that at 14-months in the APP/PS1 model of AD OPCs display a significant shrinkage resulting in a decrease in the area covered by their process domains.

**Figure 3.**
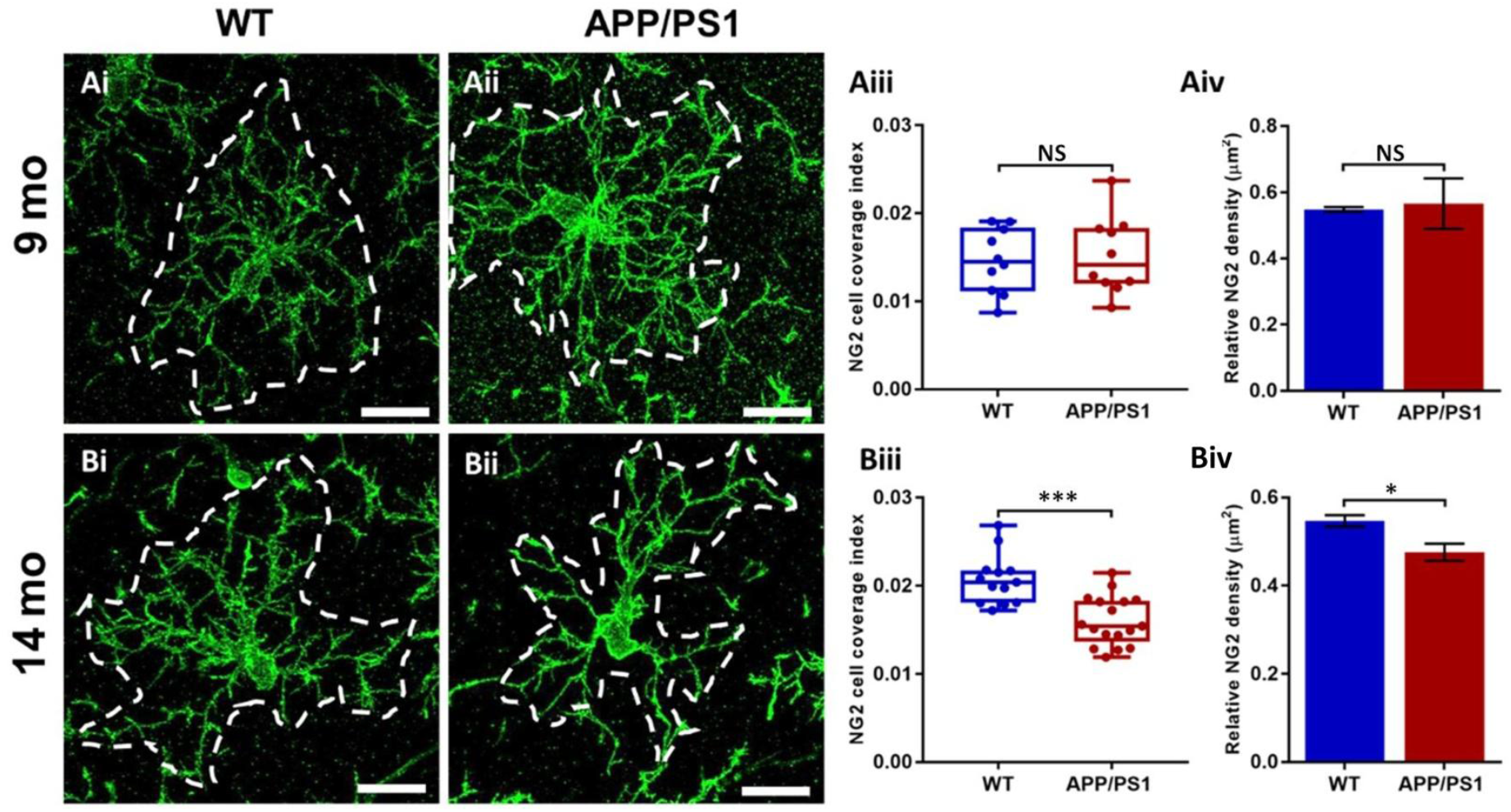
OPC process domains in the CA1 area of the hippocampus of APP/PS1 mice. Hippocampi of 9 months old and 14 months old APP/PS1 mice were examined, compared to age-matched controls, using immunofluorescence labelling for NG2 (green) to identify OPCs. High magnification confocal projections of OPCs and their process domains (indicated by broken white lines) in the 9 months old hippocampus (Ai, Aii), and the 14 months old hippocampus (Bi, Bii), in controls (Ai, Bi) and APP/PS1 (Aii, Bii). Scale bars = 20μm. (Aiii-Biii) Box-Whisker plots of the total area of OPC process domains and (Aiv, Biv) bar graphs of the coverage of the CA1 area by OPCs. Data are Mean ± SEM, \**p*<0,05; \*\*\**p*<0.001, ns= no significant; Unpaired t-test, *n*= 12 cells from 3 animals for each group.

### OPC exhibit increased process branching and cellular complexity in APP/PS1 mice

The underlying morphological changes resulting in OPC shrinkage in APP/PS1 mice were examined in further detail using Neurolucida cell tracing. Confocal images of 80-100 *z*-sections, each of 0.3μm thickness, were captured using a x100 oil objective and reconstructed and analyzed using Neurolucida 360 and Neurolucida 360 Explorer (Fig. 4A, B). Consistent with the results above, no changes in the morphological parameters of OPCs were observed between 9- and 14-months in wild-type mice (Fig. 4C-F) or in 9-month APP/PS1 OPC compared to controls (Fig. 4C-F). In contrast, OPCs were morphologically more fibrous at 14-months in APP/PS1 compared to controls, with the average number of processes per cell being unaltered (Fig. 4C), but processes displaying increased branching, with a significantly greater number of process terminals or end points (Fig. 4D; *p*<0.05, Mann Whitney test) and number of branch points or nodes (Fig. 4E; *p*<0.01, Mann Whitney test). Consequently, the Neurolucida measurement of cell complexity was 3-fold greater in 14-month APP/PS1 hippocampus compared to age-match controls (Fig. 4F; *p*<0.001, Mann Whitney test). The age-related changes in OPC complexity in APP/PS1 mice was examined further by Sholl analysis (Fig. 5A; *n*=12 cells for each group, two-way ANOVA followed by Sidak’s multiple comparisons test). As above, Sholl analysis confirmed no significant differences were detected in OPC morphology in natural aging (not illustrated). In contrast, in APP/PS1 mice there were significant differences between 9 and 14 months, with significant increases in the number of end points (Fig. 5B), the number of nodes (Fig. 5C), and in process lengths (Fig. 5D). In addition, analysis of processes length in the different branch orders identified that OPCs displayed increased process length in the distal branches (Fig. 5E). The results demonstrate OPC have increased process branching and a more fibrous appearance at 14months in APP/PS1, comparable to ‘reactive’ NG2 cells reported in human AD (Nielsen et al., 2013a) and following CNS injury (Jin et al., 2018).

**Figure 4.**
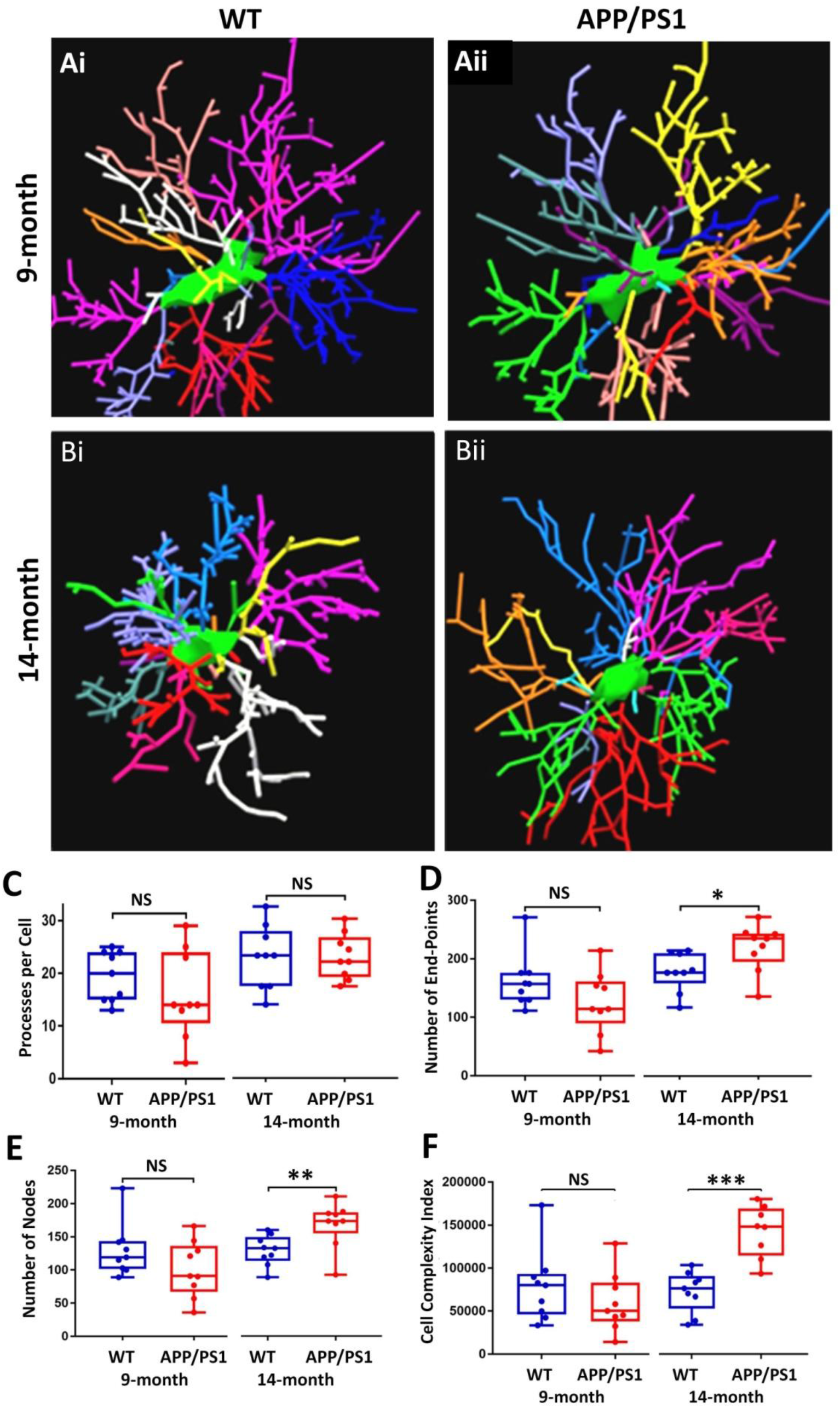
OPC morphological changes in the CA1 of the 14 months old APP/PS1 mouse model compared to an aged-matched control. Data were generated by Neurolucida 360 analysis of cells. Box-whisker plots of (A) cell body area, (B) cell body volume, (C) process volume, (D) total cell volume, (E) cell complexity, (F) ramification index. Data expressed as Mean±SEM. NS= not significant; \*\**p*<0.01; \*\*\**p*<0.001; Mann Whitney test. *n*= 9 cells from 3 animals per group.

**Figure 5.**
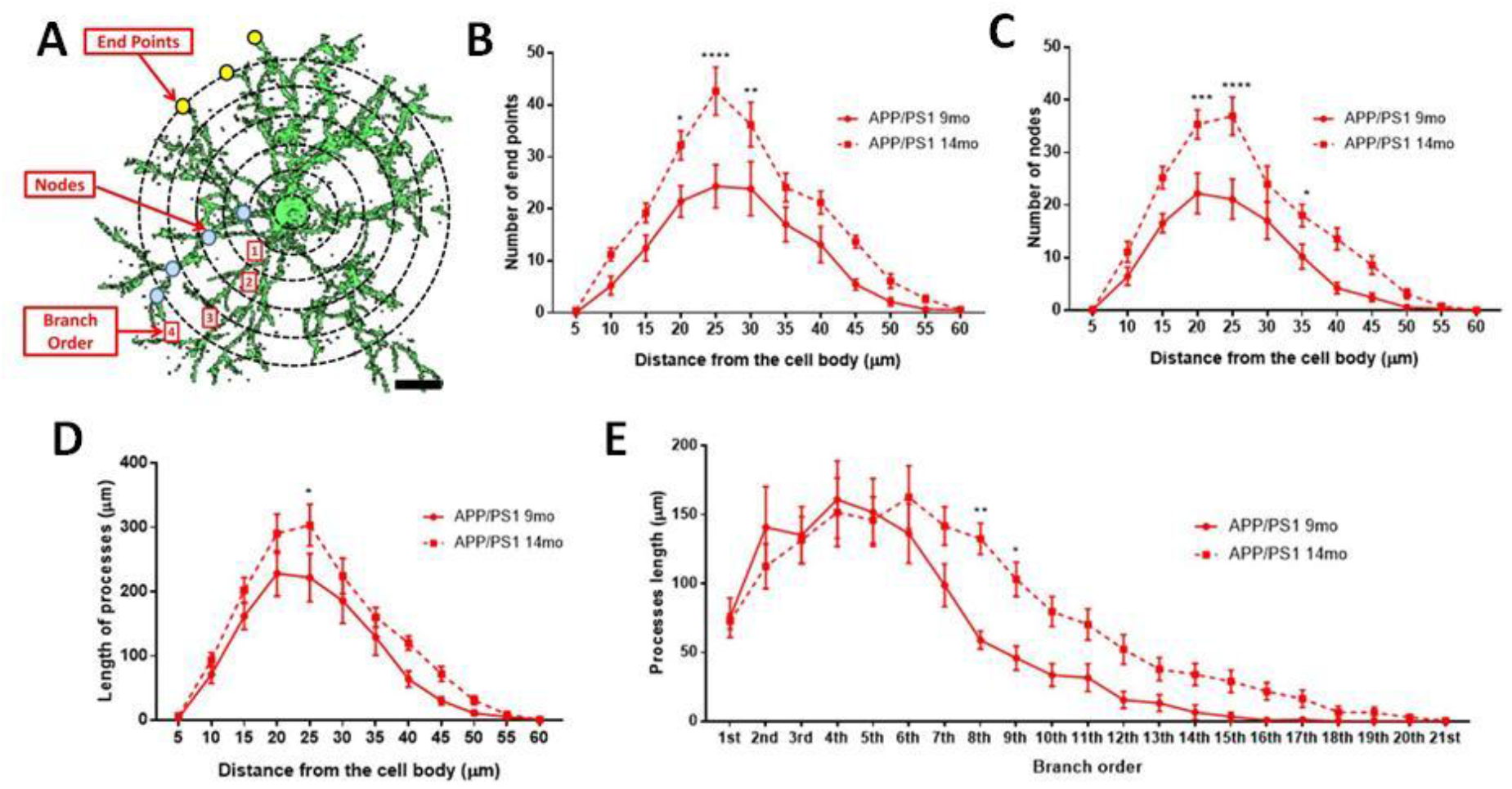
Sholl analysis of age-related changes in OPC morphology in APP/PS1. **(A)** 3D morphology of NG2 immunolabelled OPC in the CA1 area of the hippocampus (generated using isosurface rendering with Volocity software, PerkinElmer), illustrating Sholl shells (concentric circles, 5 μm apart, with the cell body in the middle), and the morphological parameters measured; the points of process branching are termed nodes (blue dots), the points where the processes intersect the Sholl shells are termed intersections (yellow dots), the st number of process terminals or end points, and the process branch order, with 1^st^ order closest to the cell body (adapted from Sholl 1953 and Rietveld et al. 2015). (**B-E**) Graphs comparing OPC morphological parameters in AAP/PS1 mice aged 9 months 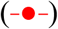 and 14 months 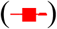; two-way ANOVA followed by Sidak’s multiple comparisons test. *p<0.05, **p<0.01, ***p<0.001, p<0.0001. *n* = 9 cells from 3 animals in each group.

## Discussion

Age-related loss of myelin has been shown to be a pathological feature of human AD (Bartzokis, 2011, Brickman et al., 2015) and in animal models of AD (Dong et al., 2018, Desai et al., 2009, Vanzulli et al., 2020). Moreover, we observed a decrease in MBP immunostaining at 14 months in the hippocampus of APP/PS1 mice, consistent with evidence that myelination is disrupted in this model of AD (Chao et al., 2018, Dong et al., 2018). The key findings of the present study are that there is a premature decrease in OPC density at 9-months, and that at 14-months OPC displayed a shrunken and fibrous morphology, indicative of morphological dystrophy. These later changes in OPCs are associated with a decline in the number of OPC sister cells, a measure of OPC self-renewal. These findings indicate that early changes in OPCs are potential predictors of myelin deficits associated with progression of AD pathology.

The generation of new myelin in the adult brain is a primary function of OPCs, which our data indicate decline in number in natural ageing between 9 and 14 months, consistent with previous studies (Young et al., 2013). Notably, this age-related loss of OPCs occurred earlier in APP/PS1, at 9 months of age, in agreement with studies indicating an early decrease in NG2 cells in AD pathology (Dong et al., 2018, Vanzulli et al., 2020). However, the number of sister cells was unaltered at 9 months in APP/PS1, suggesting the loss of OPCs at this age was not related to an overall change in self-renewal, and possibly reflects advanced OPC senescence (Zhang et al., 2019). These changes in OPCs are concomitant with a reduction in MBP immunostaining at 14-months in APP/PS1 mice compared to controls. MBP immunostaining, taken as a measure of the overall extent of myelination, was increased between 9- and 14-months in wild-type controls, but not in APP/PS1 mice, indicating hippocampal myelination is retarded in AD-like pathology (Dong et al., 2018, Desai et al., 2009, Vanzulli et al., 2020, Chao et al., 2018). We did not detect a decrease in GPR17+ or Olig2+ oligodendrocytes, suggesting the decrease in MBP immunostaining at 14-months in APP/PS1 mice is not due to a loss of oligodendrocytes per se, but may reflect a decrease in myelin production, which can involve changes in both the number and lengths of myelin sheaths (Hughes et al., 2018, Hill et al., 2018). Myelin remodelling is important for nervous system plasticity and repair (Ortiz et al., 2019, Chorghay et al., 2018, Williamson and Lyons, 2018, Foster et al., 2019), and the decline in myelination in APP/PS1 may be related to neuronal loss and learning dysfunction in these mice (Chao et al., 2018). The results provide evidence of OPC and myelin disruption in the hippocampus of APP/PS1 mice, suggesting key features of human AD are replicated in this mouse model.

Notably, the early loss of OPCs at 9-months in APP/PS1 hippocampus is followed at 14-months by atrophy of OPC process domains and a more fibrous appearance due to increased process branching. A previous study has shown changes in OPCs morphology in tissue from AD patients, in which OPCs have been reported to become less complex in the presence of amyloid-β plaques (Nielsen et al. 2013). In our study, the fibrous morphology of OPCs is similar to their reported injury response (Levine 1994). OPCs are the source of new myelinating oligodendrocytes in the adult brain (Rivers et al., 2008, Dimou et al., 2008, Zhu et al., 2008, Kang et al., 2010) and the dystrophy of OPCs in APP/PS1 mice correlates with the loss of myelin. OPC processes contact synapses in the hippocampus (Bergles et al., 2000), which regulates OPC proliferation and differentiation (Wake et al., 2011, Chen et al., 2018). The retraction of OPC processes in APP/PS1 could reflect changes in synaptic activity, which is an important feature of AD pathology. Generation of oligodendrocytes from OPC is essential for functional myelin repair (Ortiz et al., 2019), and the observed disruption of OPCs and myelination suggests these are important factors in AD.

## Conclusions

Our findings demonstrate that OPCs undergo complex age-related changes in the hippocampus of the APP/PS1 mouse model of AD-like pathology. Changes in OPCs correlate with a decline in myelination in APP/PS1. We conclude that OPC disruption is a pathological sign in AD and is a potential factor in accelerated myelin loss and cognitive decline.

## Declarations

### Ethics approval and consent to participate

All animal procedures were carried out in accordance with the Animals (Scientific Procedures) Act 1986 of the UK.

### Consent for publication

Not Applicable.

### Availability of data and material

All data generated or analysed during this study are included in this published article.

### Authors’ contributions

IC-D-L-R: Formal Analysis; Investigation; Methodology; Writing - original draft.

GF: Formal Analysis; Investigation; Methodology; Validation.

ADR: Investigation.

AV: Conceptualiztion; Writing - review & editing.

DG-N: Conceptualization; Data curation; Formal analysis; Funding acquisition;

Project administration; Resources; Supervision; Validation; Visualization; Writing - review & editing.

AMB: Conceptualization; Data curation; Formal analysis; Funding acquisition;

Project administration; Resources; Supervision; Validation; Visualization; Writing - original draft; Writing - review & editing.

## Acknowledgements

Not Applicable

